# Stony Coral Symbioses Show Variable Responses to Future Ocean Conditions

**DOI:** 10.64898/2026.03.06.710183

**Authors:** Mariana Rocha de Souza, Janaya Bruce, Annick Cros, Christopher P. Jury, Crawford Drury, Robert J. Toonen

**Affiliations:** Hawai’i Institute of Marine Biology, School of Ocean and Earth Science and Technology, University of Hawai’i at Mānoa, Honolulu, HI 96744, USA; University of California, Santa Cruz, CA 95064, USA; The Nature Conservancy, CA, USA

## Abstract

Coral reefs support over a quarter of marine species and nearly a billion people worldwide but are also among the ecosystems most threatened by anthropogenic impacts. There is long-standing debate about whether coral symbioses will be disrupted or respond adaptively under future ocean conditions. Using a factorial 2.5-year future-ocean mesocosm experiment across eight coral species representing the major coral lineages, we tracked symbiont community shifts within replicate fragments from the same individual coral. Some corals exhibited stochastic divergence consistent with dysbiosis, whereas others showed deterministic, thermally adaptive shifts. Heat stress generally reduced symbiont diversity and promoted predictable restructuring, supporting deterministic processes under moderate stress but stochastic dysbiosis under extreme conditions. We propose that adaptive and stochastic responses represent endpoints along a continuum of host-orchestrated symbiont sorting. This study bridges coral reef ecology with broader host–microbiome theory, offering an integrated perspective on how symbiotic systems may respond to environmental change.

**Teaser:** Long-term experiments reveal that corals follow distinct paths of symbiont change under heat stress, from adaptation to dysbiosis

## Introduction

Mass coral bleaching events are a recent phenomenon (*1*) directly attributed to heat stress caused by increasing ocean temperatures (*2, 3*). The frequency and severity of thermal stress has increased significantly in recent decades, including record-breaking sea surface temperatures in July 2023 with the past 10 years being the warmest decade in recorded history (*4*). These conditions have caused the near-extinction of

Caribbean *Acropora* corals in Florida (*5*) and raised concerns that coral reefs may have reached a global tipping point (*6*). Among the many stressors on coral reefs, ocean warming is among the most certain to persist and intensify, leading to substantial increases in global ocean temperatures even if greenhouse gas emissions ceased today (*7*). As part of this CO_2_ emissions trajectory, ocean chemistry is also changing, creating conditions that are energetically unfavorable for coral calcification and reproduction. Many studies predict that the combination of ocean acidification and warming will lead to the functional collapse of coral reef ecosystems over the next few decades, driving significant losses in biodiversity, ecosystem services, and ultimately the dissolution of the carbonate structure which creates reef ecosystems (*8*–*11*).

Symbiotic dinoflagellates have a foundational influence on the climate-related phenotypes of individual coral holobionts, which dictate the long-term persistence of coral reef ecosystems. There are eleven genera within Symbiodiniaceae (*12, 13*) and eight named species of Symbiodiniaceae with life histories that range from free-living to obligate symbionts of reef-building corals and other marine invertebrates (*14*).

Symbiodiniaceae species identity and community composition greatly affects host performance (*15*–*17*) and impacts the ability of the holobiont to thrive at different latitudes, depths, irradiances, and temperatures (*15, 18*–*20*). Some endosymbiont species increase heat tolerance in corals (*15, 16, 21*), while other species are opportunistic and increase susceptibility of corals to disease (*22*), or might even be considered parasitic in some conditions (*23*–*25*).

Despite the high diversity of Symbiodiniaceae, host-symbiont associations are not random and exhibit varying degrees of specificity. Some coral species associate with only a single dominant or a few co-dominant symbiont types (*26, 27*), while others associate with many different symbiont types (*28*). The degree of specificity can depend on the host, the symbiont, or the environment (*28, 29*). Most studies have focused on the contribution of individual symbiont taxa to the holobiont environmental stress response, an approach called the ‘selection effect’ which assumes that the independent function of the dominant species drives symbiont community function (*30*–*32*). However, overall symbiont community composition may also influence holobiont stress tolerance via the ‘complementarity effect’ (*30, 33*) an emergent property of community diversity in which the function of symbiont taxa is dependent on the interactions arising from their community context, including niche partitioning and other positive interactions within an individual coral holobiont (*34*).

Understanding the temporal dynamics of coral symbiont community composition in response to environmental stressors is essential to both the basic biology and management of coral reef systems. Our study aims to elucidate the nature of symbiont changes to test two different opposing hypotheses currently in the literature. The Anna Karenina Principle (AKP) for animal microbiomes (*35*) derives its name from the famous opening line of Tolstoy’s novel, Anna Karenina: “all happy families are all alike; each unhappy family is unhappy in its own way.” When applied as a hypothesis to host-associated microbiomes, AKP posits that individual stressors do not exert consistent, deterministic influences on community composition. Instead, stressors introduce stochastic variations that disrupt species interactions, causing stressed microbiota to break down unpredictably and exhibit unique patterns of dysbiosis. Patterns consistent with AKP have been observed in a wide range of organisms including humans (*23*–*25*). To date, approximately 50% of diseases in humans follow the AKP Principle, while the remaining diseases exhibited either non-AKP effects (25%) or effects unrelated to AKP (*36*).

Alternatively, the Adaptive Bleaching Hypothesis (ABH) posits that loss of endosymbionts during coral bleaching provides the niche space required for corals to reconstitute the community composition of their algal symbionts in response to environmental stress (*37*–*39*). Some corals clearly change the relative proportion of tolerant endosymbionts hosted in response to thermal stress (*40*–*43*), while others maintain stable associations despite bleaching, or revert to pre-bleaching symbiont composition when recovering from stress(*44*–*46*). A higher proportion of heat-tolerant symbionts can be found in corals at sites with variable microhabitats and warmer average temperatures (*47*–*49*); however, there are prominent counterexamples where corals do not change their symbiont community after bleaching (*50, 51*). The Adaptive Bleaching Hypothesis remains controversial, in part due to the diversity of symbiont transmission modes (maternally or environmentally acquired) and specificity dynamics that underlie the continuum of coral-algal relationships (*52*–*56*), and in part because it is difficult to envision a mechanism by which selection favors a holobiont as an integrated individual as opposed to a multi-species community (*57, 58*). However, recent theoretical work shows that coral host-mediated species selection of symbionts followed by microbial competition can drive host-symbiont integration consistent with the ABH (*59, 59, 60*).

A key element of the AKP is the bidirectional interaction between the coral immune system and the core microbiome. The notion that stability is associated with a healthy microbiome or host physiology mirrors the notion that dysbiosis (or increasing heterogeneity) is often associated with the diseased microbiome and abnormal host physiology which can lead to host mortality. Non-AKP dynamics indicate that stressed organisms either maintain stability of their microbial community or change it in a deterministic manner that supports or improves fitness during environmental challenges. Such a responsive shift in the microbiome underscores an adaptive relationship between hosts and their associated microorganisms, suggesting the capacity for selection to increase holobiont fitness in the face of stressors. Thus, stochastic increases in symbiont diversity are predicted to be unstable due to colonization by selfish microbes that do not benefit the host, whereas deterministic selection of beneficial symbionts by the host favors long-term stability of the holobiont association (*60*).

Here we investigate patterns of coral holobiont community change under future ocean conditions to determine whether they follow stochastic patterns (AKP) or show directional trends towards symbionts (ABH) that have the potential to increase holobiont survivorship or fitness. To quantitatively distinguish between stochastic and deterministic symbiont responses, we calculated the difference in Hill numbers (Δq) between each treatment and control group across diversity orders (q = 0–3), where positive values indicate increased stochasticity (supporting the Anna Karenina Principle) and negative values reflect directional change consistent with the Adaptive Bleaching Hypothesis. We exposed 8 species of corals from 6 locations with diverse environmental histories to ∼2.5 years of a factorial array of simulated future climate conditions (ambient, acidified, heated, dual stressor) in experimental mesocosms (*11, 61*) and evaluated changes in symbiont community using ITS2 sequencing. Quantifying the nature of these responses is central to understanding the diversity of host-microorganism interactions and their implications for coral reefs under future ocean conditions.

## Results

### Coral collection and experimental treatments

Corals were collected from 6 locations with differing environmental histories and chemical-physical properties around O’ahu, Hawai’i (**Fig. 1a, Supplementary Table 1**) and maintained in present-day average, acidified (-0.2 pH units below present-day), heated (+2 °C above present-day), or dual-stressor future ocean conditions (-0.2 pH, +2 °C relative to present-day). These species collectively account for >95% of the coral cover on Hawaiian reefs (*62, 63*), and provide representatives of both major evolutionary lineages of scleractinians (Complexa and Robusta), three of the primary reef-building coral families (Acroporidae, Pocilloporidae, and Poritidae), and all four of the major life history strategies ((*64*), including competitive (*Montipora capitata, Pocillopora meandrina*, and *Porites compressa*), generalist (*Montipora flabellata* and *Montipora patula*), stress-tolerant (*Porites lobata* and *Porites evermanni*), and weedy strategies (*Pocillopora acuta*), exhibited by corals (*65*).

**Figure 1.**
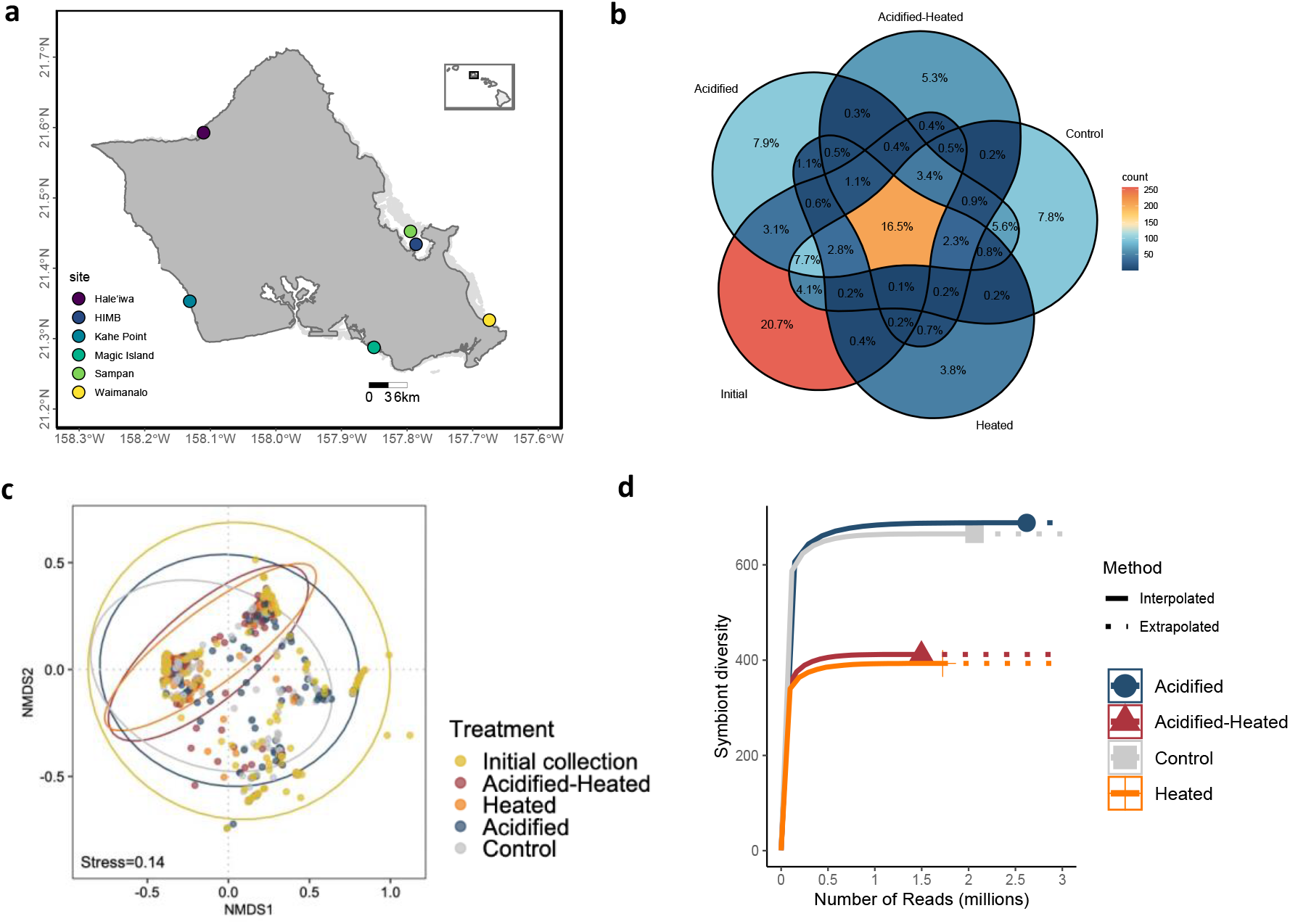
**a)** Map of O’ahu with the sites of the original collections. This broad spatial sampling of corals helped to ensure that a representative sample of the genetic variation present in these species from O‘ahu was included in the study. The collection sites were chosen to represent a range of environmental conditions around O‘ahu, serving as a natural laboratory (**Supplementary Table 1**); **b)**Venn diagram of the number of algal symbionts found in the corals in the experiment in each treatment; **c)** NMDS of the symbiont composition in each treatment; **d)** Chao1 coral symbiont species richness estimates for each treatment.

The heated treatments were above the nominal bleaching threshold for 3.5 months per year, during the warmest season resulting in ∼24 Degree Heating Weeks (DHW) accumulated annually (*11*), which is slightly more than the ∼22 DHW accumulated in the Florida Keys during the unprecedented 2023 marine heat wave (NOAA Coral Watch). The acidified treatment resulted in a pH decline of about 0.2 below present-day values, and about 0.3 below preindustrial levels. When combined with the heat treatment, this condition has been suggested in the literature to cause reefs to shift from net calcification to net carbonate dissolution (*9, 66*), yet we saw about 65% survivorship of corals, which is much higher than generally projected (*11*).

### Symbiodiniaceae community composition changes among treatments

In total, 904 of the initial 1184 coral fragments (76.4%) survived the ∼2.5-year experiment. A subset of the survivors in the Control and Ocean Acidification treatments died prior to sampling due to a hurricane, which disrupted seawater flow to a subset of the Control and Ocean Acidification mesocosms. From those 148 samples which survived the Acidified Heated double stressors, 147 survived the Heated condition, 206 survived the Acidified condition and 171 survived the control condition (**Supplementary Table 2**). A total of 1246 Symbiodiniaceae types (including the post-treatment samples and the initial state for each colony collected from the field; see Methods) were identified using SymPortal (*67*); 69% belonged to the genus *Cladocopium*, 20% belonged to the genus *Durusdinium* and the remaining belonged to the genera *Symbiodinium* (7%), *Breviolum* (3%), *Fugacium* (0.4%), and *Gerakladium* (0.2%).

Using the full diversity of sequence variants identified through SymPortal, 16.5% of the symbiont diversity was shared among all treatments, while 3.8% of symbionts were only present in the Heated treatment, and 5.3% were unique to the dual stressor treatment (**Figure 1b**). Algal symbiont community differed significantly among treatments (PERMANOVA, F_4_=5.816, p=0.001; **Figure 1c, Supplementary Table 3**). Pairwise comparisons revealed that the two treatments with temperature stress (Heated and Acidified-heated) were significantly different from the other treatment conditions, whereas acidified treatment did not significantly differ from control or initial collection (**Figure 1c, Supplementary Table 3**).

Corals grown under elevated temperature treatments (Acidified-Heated and Heated) presented reduced Symbiodiniaceae diversity relative to corals in the other treatments (Chao 1; **Figure 1d**), emphasizing the negative impact of temperature stress on diversity in a complex landscape of differential effects of environmental stress.

To identify the symbionts most responsible for differences among treatments and assess the extent of community shifts from the control, we created biplots of the symbiont communities in corals exposed to each stress treatment, alongside those from the control group. (**Figure 2a-c, Supplementary Table 4**). In all treatments, heat-resilient symbionts from the genus *Durusdinium* (D4, D1, D6, D1dg) clustered together, while various types of *Cladocopium* occupied a distinct region of the biplot. Coral symbionts exposed to the heat stress treatments (Acidified-Heated and Heated) showed similarities (**Figure 2a, b**), with most corals in the heat stress treatment (**Figure 2b**) clustering near *Durusdinium* and C15, Symbiodiniaceae taxa known for their heat stress resilience (*15, 16, 68*). Symbionts from corals exposed to acidified conditions were similar to the symbionts from the control treatment across the same multidimensional space (**Figure 2c**), indicating similar symbiont composition. Loadings were higher in corals exposed to temperature stress when compared to Acidified conditions (**Supplementary Table 4**). These results highlight that temperature changes, rather than acidification, was the primary driver of differentiation in symbiont composition.

**Figure 2.**
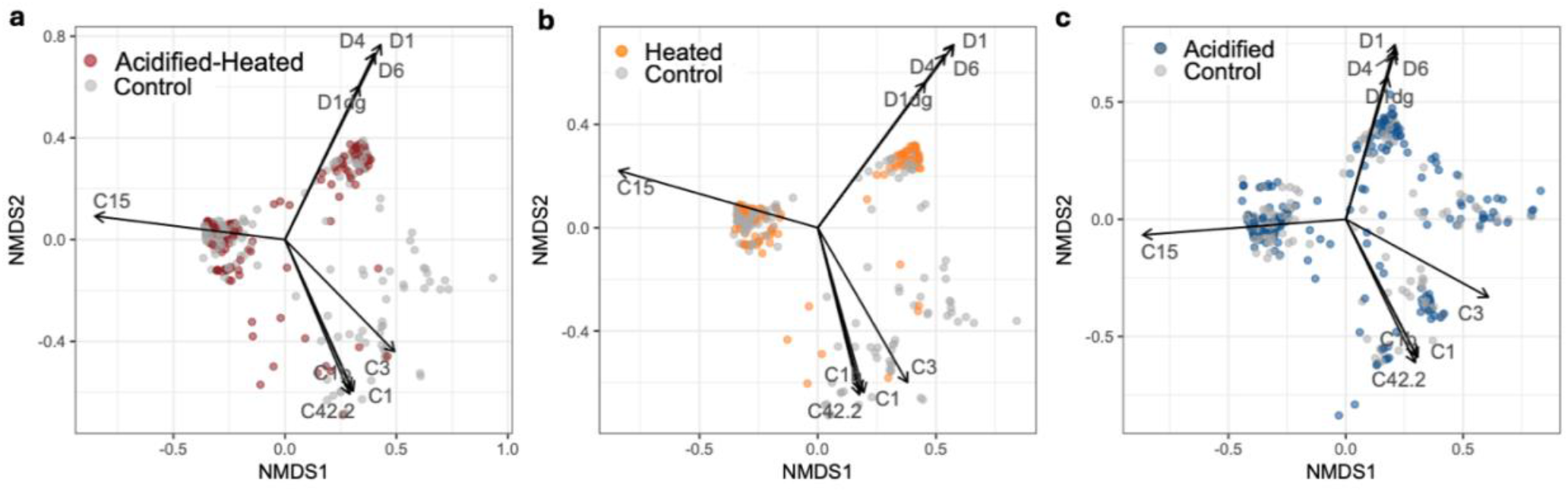
Biplot showing the relationship between symbionts present in corals exposed to each treatment in the experiment compared to control treatment. **a)** Biplot of coral under Acidified-Heated and Control; **b)** Biplot of corals under Heated and Control; **c)** Biplot of corals under Acidified and Control. Loadings are reported in **Supplementary Table 4**.

### Effects of ABH and AKP across coral species, collection sites, and treatments

Using ITS2-derived symbiont community composition at the end of the experiment for both control and treatment conditions, we calculated the difference in Hill number (q) between each treatment and control (Δq). The Hill numbers present a “diversity profile”, which offers a series of diversity measures corresponding to different diversity order (q = 0, 1, 2, 3), weighted differently by species abundances (*35, 36*). Lower orders (e.g., *q* = 0) count all species equally, regardless of their abundance, whereas higher orders place progressively greater weight on common or dominant species. To account for the symbiont taxa present, as well as the relative abundance of each taxon, we used q=2, representing the Simpson Diversity Index, which increases the weight of the dominant species. Positive Δq values represent stochasticity consistent with the Anna Karenina Principle (AKP), while negative Δq values represent deterministic change consistent with the Adaptive Bleaching Hypothesis (ABH).

When examining site-specific Δq (**Figure 3a**), we observe large differences in Δq within the same species across locations under most treatments. For example, under Acidified-Heated, *Porites evermanni* exhibits positive Δq in all locations, except for Magic Island. In contrast, in heated conditions, most *P. evermanni* samples present negative Δq, with the exceptions of Waimānalo and Sampan channel. Some species, such as *Montipora flabellata* predominantly display negative Δq across most locations.

**Figure 3.**
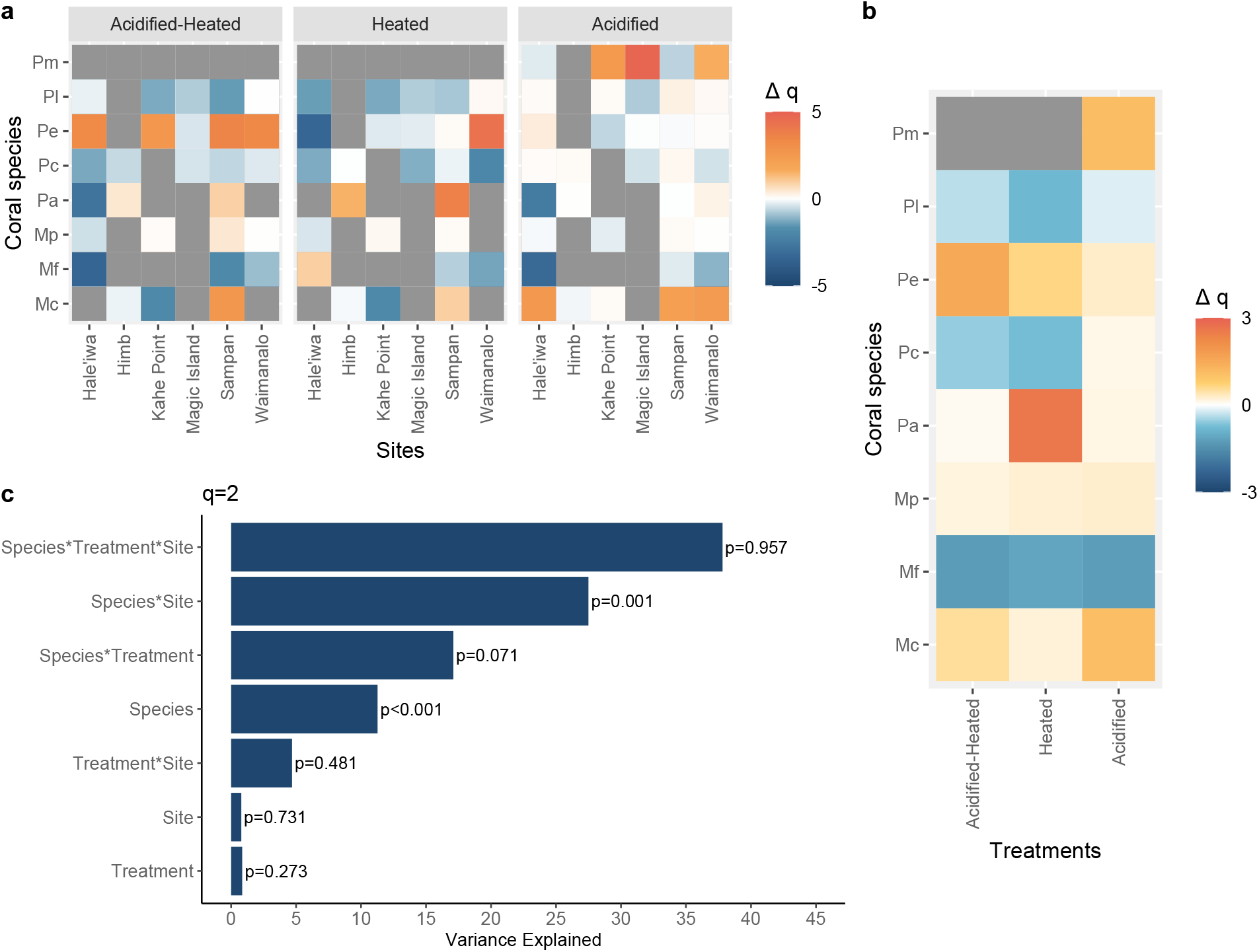
Heatmap of the change in Hill number (Δq) for **a)** each of the eight species considering the different sites of origin, **b)** all origin sites combined exposed to high temperature with high pCO_2_ (acidified-heated), high temperature with ambient pCO_2_ (heated), and high pCO_2_ with ambient temperature (acidified) treatments and **c)** Variance explained of each one of the variables used in the study. Δ q is the difference in Hill number for q2 for fragments of each species exposed to the treatments compared to the Hill number of the same individual exposed to the control treatment, where a positive mean Δq indicates AKP and negative mean Δq indicates ABH. Coral species abbreviations are as follows: Mc= *Montipora capitata*, Mf= *Montipora flabellata* and Mp= *Montipora patula*, Pm= *Pocillopora meandrina*, Pc= *Porites compressa*, Pl= *Porites lobata*, Pe= *Porites evermanni*, Pa= *Pocillopora acuta*. Gray cells indicate instances where no samples from a species at a specific location survived the treatment.

Interestingly, under the Acidified-Heated and Heated treatments, 61% and 63% of the colonies, respectively, exhibited negative Δq (ABH). In contrast, under the acidified treatment, approximately 75% of the samples had Δq values close to zero (within the range of -0.9 to 0.9). This increase in the proportion of Δq values near zero suggests a comparatively smaller difference in symbiont community composition between coral samples maintained under acidified conditions and those in the control treatment. These findings indicate that acidification had a smaller effect on symbiont community composition than temperature, leading to very small difference in symbiont community diversity compared to control, whereas thermal stress lead to symbiont change in a deterministic way (ABH).

The mean Δq per species (**Figure 3b)** shows that certain species, such as *Porites lobata* and *Montipora flabellata*, consistently exhibit negative mean Δq values (consistent with ABH) across all treatments. In contrast, *Montipora capitata, Montipora patula*, and *Porites evermanni* display positive mean Δq values (consistent with AKP) in all treatments. Similar to the observations in **Figure 3a**, most samples under the acidified treatment had positive Δq values, often smaller and closer to zero. This pattern aligns with the AKP model, suggesting that symbiont community differences were minimal and largely stochastic when compared to clonal fragments of the same individuals maintained under ambient conditions.

The analysis of the deviance shows that all variables in the experiment — treatment, site of origin, and species— make significant contributions (p<0.005) to symbiont community composition (**Supplementary Table 5)**. In terms of explained variance, species and the interaction of species and site were the most significant factor driving the symbiont community change (**Figure 3c, Supplementary Table 6**). This interaction highlights the role of biological differences among coral species affecting the symbiont community and how those differences interact with the environmental conditions to which the coral holobiont is exposed at each site.

Among surviving corals, the proportions following AKP and ABH strategies did not differ significantly across treatments (p = 0.16; **Supplementary Figure 1**).

## Discussion

The effect of heat stress on corals, leading to a loss of symbionts and resulting in coral bleaching, has been documented in many studies (*1, 69, 70*). In contrast, the effect of acidification on coral symbiont density and bleaching outcomes have generated contrasting results (*71*–*75*). Here we examine changes to the symbiont community composition among clonal ramets of corals maintained in long-term coral reef mesocosms maintained for ∼2.5 years under a factorial design of acidification and warming predicted in the year 2075 under a high CO_2_ emission scenario (*11*). We use eight major reef building coral species (collectively accounting for >95% of coral cover in Hawai‘i) that covers a breadth of the taxonomic and functional diversity of reefs from throughout the Central Pacific. We find no significant effect of acidification treatments on Symbiodiniaceae community structure and heat stress as the main driver of Symbiodiniaceae community composition change among experimental treatments. Both heated and heated-acidified treatments show a similar shift in Symbiodiniaceae community composition relative to initial collections and control treatments. In contrast, fragments maintained under control and acidified-only treatments were not significantly different from one another. In fact, across a diverse assemblage of corals exposed to future ocean conditions for roughly 2.5 years, we find that the effects of ocean acidification on the coral-algal symbiosis are undetectable relative to control.

Coral responses to thermal stress vary by both host and symbiont species, and among locations, habitat and environmental conditions (*20, 76, 77*) but the underlying mechanisms and role of the symbiotic community in driving such variability remains less understood. For example, de Souza et al. (*78*), reported that the algal symbiont composition of *Montipora capitata* corals varied across hydrodynamic regimes in Kāne’ohe Bay under different environmental conditions, including depth, temperature regimes, and sedimentation. Variations in symbiont community composition corresponded to local environmental factors, with depth and temperature variability being the strongest drivers across sites. Conversely, (*79*) showed that corals on the hottest reefs in the Persian Gulf maintain associations with the same symbionts across 1.5 years despite extreme seasonal temperature changes and acute heat stress (≥35°C). Subsequently,(*46*) found that the overall algal symbiont community composition was also consistent and largely unchanged among sites, despite a significant increase in the relative abundance of the heat-tolerant symbiont *Durusdinium* during a mass bleaching event, indicating that site history is a key factor influencing Symbiodiniaceae community composition in *M. capitata* corals within Kāne‘ohe Bay. Building on this previous work, here we find that the dominant predictors of coral symbiont community composition remain the species-specific associations and provenance of the individual fragments. We find symbiont community composition is driven primarily by the species of the coral host and that provenance provides greater predictive power than treatment exposure for symbiont community composition, although treatment is also significant. Our findings are consistent with recent field surveys showing that among 600 tagged colonies, fine-scale environmental factors such as depth and temperature variability overwhelmed an increase in the heat-tolerant symbiont *Durusdinium* during a natural bleaching event and that site explained most of the observed variance in symbiont community composition (*46*).

Overall, we found that experimental treatment had significantly less effect on the symbiont community composition than coral host species and collection location, and among treatments ocean warming had a much greater effect than acidification. Ocean acidification led to minimal and largely stochastic symbiont community change (AKP) while temperature decreased the symbiont diversity and drove the symbiont community in a deterministic way more consistent with ABH. Coral species from the same genus respond differently to the treatment conditions, and those responses can be modified by legacy effects from the site of collection. These species- and site-specific responses may reflect legacy effects from the site of collection, indicating that corals exposed to long term environmental conditions might acclimate to local conditions, creating an “environmental memory” (*46, 80*). This memory, via physiological or symbiont composition changes, influences coral bleaching thresholds and survivorship under later stress (*81*).

We also found that thermal stress changes the community composition loadings of the biplots (**Table S4**), but these changes do not completely replace the underlying species and location patterns. This helps explain the varying results seen in experiments with different coral species. Prior work found that coral algal symbiont beta diversity increased (AKP effect) during exposure to thermal stress (*32*) in *Acropora millepora*. Contrastingly, we found that heat stress led to an overall decrease in algal symbiont beta diversity (ABH) relative to the same fragments maintained under control conditions, while acidification made little difference to symbiont diversity compared to control, but with clear differences among species that make further generalizations difficult.

Symbiont selection, as described by the Adaptive Bleaching Hypothesis (ABH), may enhance coral survivorship (*37, 38*), highlighting the fitness advantages for corals of reducing symbiont diversity via a directional shift to stress-resilient symbionts. Consequently, corals equipped with such selected symbionts might demonstrate enhanced resilience to stress factors, mitigating bleaching and mortality risks (e.g. Cunning (*39*)). Survivorship varied among species: no *Pocillopora meandrina* fragments survived the high temperature treatments (either the acidified-heated or heated), and within coral species, individuals had varying levels of mortality depending on the site of origin. Unfortunately, it is not possible to test the correlation of survivorship and ABH in our samples, because symbiont communities could not be identified at the endpoints for samples that died during the exposure, so it is impossible to know whether they followed ABH or AKP. When comparing only survivors, the number of surviving samples following ABH is slightly greater than those following AKP, but that difference is not significant (**Supplementary Figure 2**). Unfortunately, there is no way to compare the strategies of corals that did not survive the experiment without intensive temporal sampling, which we did not perform.

Coral species and the interaction of species and the site of origin are the greatest predictors of change in the symbiont community composition (**Supplementary Table 6**). For example, *Porites lobata* from Magic Island and Waimānalo; *Montipora patula* from Kahe Point and Sampan; and *Montipora flabelata* from Sampan, Waimānalo and Hale‘iwa show changes in symbiont community consistent with AKP whereas individuals of the same species originating from other sites do not. This contrasts with *Montipora capitata* and *Porites compressa* which follow AKP under acidified conditions only, and *Pocillopora acuta* that conforms to AKP under all treatments from all sites except for samples collected from Hale‘iwa. Among collection sites, both HIMB and Magic Island have exceptional environmental conditions. Kāne‘ohe Bay corals (where HIMB and Sampan site is located) experience high temperature and pCO_2_ conditions that are not expected to occur regularly at more oceanic sites around O‘ahu until late century (*65, 82, 83*), whereas the reefs around Magic Island are exposed to frequent rain-driven effluent and increased nutrients from the Ala Wai Canal (*84*).

Given the number of samples, species, treatments and the multiyear length of the experiment, our study provides an incredible opportunity to investigate evidence for the two primary hypotheses for coral holobiont change under environmental stress: the Anna Karenina Principle (AKP) and the Adaptive Bleaching Hypothesis (ABH). We use this experiment to test the importance of coral host species, site of initial collection and future ocean conditions (warming and acidification) in influencing the symbiont composition of coral holobionts. We find evidence that both processes occur, and our results support the latest theory on holobiont evolution that predict conditions under which host-orchestrated species sorting should increase or decrease symbiont diversity. Roughgarden (*60*) develops a mathematical model for the population dynamics of holobionts that frames a horizontally transmitted microbiome as a system of collective inheritance, in which the environmental microbial source pool plays a role analogous to the gamete pool in sexual reproduction. In each generation, hosts acquire their microbiome by *Poisson sampling* from this environmental pool, and the composition of that pool is shaped by the combined output of all hosts in the population. A central insight of the model is that microbes beneficial to the host may incur a cost to their own within-host growth rate, allowing them to be displaced by otherwise identical “freeloader” strains that do not contribute to holobiont fitness. To maintain a functional partnership, hosts must therefore actively filter incoming microbes **—** a process termed host-orchestrated species sorting (HOSS) — and allow only certain taxa to establish. Once inside the host, competition among these allowed microbes further refines the community. This mechanism, rather than coevolution or multilevel selection, is predicted to be the main driver of stable microbiome–host integration.

Site-specific differences in community composition in our experiment are consistent with HOSS followed by microbial competition driven by local environmental conditions. We observe the greatest overall spread in holobiont community composition among the initial O‘ahu-wide collection, which becomes reduced when individuals are maintained in a common garden (control treatment) and reduced further under thermal stress (**Figure 1c**). This pattern is consistent with competitive adjustment of the microbial community in response to environmental manipulations. In Roughgarden’s framework, stringent HOSS or strong microbial competitive exclusion reduces symbiont richness — an outcome consistent with the Adaptive Bleaching Hypothesis (ABH) as an adaptive response to holobiont selection. Loss of HOSS control, by contrast, allows colonization by non-mutualistic or novel taxa, increasing diversity in a manner consistent with the Anna Karenina Principle (AKP). Our data show both patterns: warming treatments decrease symbiont diversity in line with ABH, while in other cases we detect AKP-like increases, including symbionts (*Breviolum, Fugacium*, and *Gerakladium*) not previously observed in healthy Hawaiian corals, potentially signaling loss of HOSS control and the onset of dysbiosis. We therefore view AKP and ABH as ends of a mechanistic continuum of host-orchestrated control of the coral symbiotic community, consistent with the HOSS model of holobiont evolution.

## Matherial and Methods

### Experimental Design

The experiment was conducted in a 48-tank outdoor, flow-through mesocosm system at the Hawai’i Institute of Marine Biology (HIMB) on Moku o Lo‘e (Coconut Island), in Kāne‘ohe Bay on the island of O‘ahu in the Hawaiian archipelago. Tank and experiment set up are detailed elsewhere (*11, 61, 83, 85, 86*). In summary, unfiltered seawater pumped continuously and directly from the adjacent coral reef slope fed the fully factorial design with a total of four treatments consisting of present-day pH and temperature (control), present-day temperature with acidification (acidified treatment: -0.2 pH units relative to control), present-day pH with elevated temperature (heated treatment: +2 °C relative to control), or both acidification and elevated temperature (combined future ocean treatment: -0.2 pH units and +2 °C relative to control), with 10 replicate mesocosms per treatment. Corals were collected at 2±1 m depth at 6 different locations around O‘ahu (Waimānalo, Hale‘iwa, Sampan Channel, Kahe Point, Magic Island and from the reefs surrounding HIMB **—** Moku o Lo‘e ; **Figure 1**) to help ensure that a representative sample of their phenotypic and genotypic diversity was included in the study. The collection sites were chosen to represent a range of environmental conditions around O’ahu (*83*), serving as a natural laboratory (**Supplementary Table 1**).

### Initial Collection

DNA was extracted using the E.Z.N.A. Tissue DNA kit (Omega Bio-tek, Norcross GA) from *Montipora capitata* (22 colonies), *Mon. flabellata* (14 colonies), *Mon. patula* (23 colonies), *Porites compressa* (27 colonies), *Por. evermanni* (28 colonies), *Por. lobata* (29 colonies), *Pocillopora acuta* (24 colonies), and *Poc. meandrina* (23 colonies), for a total of 190 samples (**Supplementary Table 2**). From each colony, a 2 mm^2^ piece of coral was ground and incubated overnight at 55 ^°^ C in 200 µl of lysis buffer and 25 µl of proteinase K. DNA was eluted once in 50 µL and twice in 100 µL of elution buffer following manufacturer protocols. Extracts were visualized on an agarose gel, and the cleanest elution was used for PCR amplification. Each parent colony was genotyped using available microsatellite loci (*87, 88*) to ensure that they were distinct genets, and our results were not biased by inclusion of multiple clonally derived colonies. Amplicons were generated via PCR using microsatellite primers tagged with unique barcodes, pooled equimolarly, and sequenced on an Illumina MiSeq platform (v3 2x300 PE) at the Hawai‘i Institute of Marine Biology following previously published methods (*89*). Alleles were called using a custom bioinformatic genotyping workflow, which were then converted to GenoDive v. 2.0b27 (*90*) file format for analyses as outlined in (*89*). Using the ‘assign clones’ feature of GenoDive, we tested whether coral colonies sampled in the field had a unique multilocus genotype. To be conservative, we allowed for up to two scoring errors among individuals and then manually checked each potential clone against the location of collection.

### Final Collection

Replicate clonal fragments (ramets) of each genetically unique coral colony (genet) were included in all four treatments. Most colonies contributed 1 nubbin per treatment. Coral samples who survived the treatment were collected on November 19^th^ and 20th, 2018, about 2.5 years after they were first placed in the treatment tanks. Tissue biopsies were preserved in SSD buffer (*91*)until they could be extracted. The E.Z.N.A. Tissue DNA Kit (Omega-Biotek, Norcross, GA) was used to extract genomic DNA from each of 60 ramets of *Mon. capitata*, 77 of *Montipora flabellata*, 84 of *Montipora patula*, 115 of *Porites compressa*, 106 of *Porites lobata*, 102 of *Porites evermanni*, 50 of *Pocillopora acuta*, and 34 of *Pocillopora meandrina*, for a total of 628 fragments extracted and sequenced.

#### Symbiodiniaceae ITS2 Amplicon Sequencing Library Preparation

Coral symbiont ITS2 amplicon library preparation and sequencing followed that previously reported by de Souza et al. (*78*). Briefly, ITS2 was amplified for each sample using individually barcoded primers, that were pooled equimolarly and sequenced on the Illumina MiSeq platform (v3 2x300bp PE). Raw sequences were demultiplexed using Cutadapt (*92*). Demultiplexed forward and reverse fastq. gz files were remotely submitted to SymPortal (*67*) to predict putative Symbiodiniaceae taxa (*14*) from the ITS2 marker sequences. Sequence quality control was conducted as part of the SymPortal pipeline using Mothur 1.39.5(*93*), the BLAST+ suite of executables (*94*) and minimum entropy decomposition (*95*). Here we restrict our use to ‘Symbiodiniaceae type’ A type refers to Symbiodiniaceae taxa that have a specific sequence as their most abundant sequence.

### Statistical Analysis

All analyses were done in the R statistical environment (*96*). To account for the effects of moving corals into experimental tanks, analysis of colonies at the time of initial collection, the control treatment (unfiltered Kāne‘ohe Bay water in the tanks), and each of the remaining heated and acidified treatments were done separately. For this study, statistics were performed using ITS2 sequence (ASV) data.

Hill numbers were calculated using the *hillR* package. Pairwise beta-diversity, at the diversity order q=2, representing species richness, was calculated for all samples within the control treatment and for each one of the treatments, respectively. Delta q (difference in Hill number from corals exposed to the treatment to control) were also calculated in R.

General linear model was calculated in R using the function *glm* and *car::Anova*. To evaluate the importance of the variables, we looked at the variance explained for each variable by calculating the hierarchical partitioning in R using the package *glmm*.*hp*.

Bray-Curtis dissimilarity of relative abundance of the Symbiodiniaceae community composition for each species was tested by permutational multivariate analysis of variance (PERMANOVA) in the function *adonis* (for effect of species), and pairwise.*adonis* (for pairwise PERMANOVA) in the *vegan* package (*97*) each with 999 permutations. The function *metaMDS* was used in the R package *vegan* (*97*) to generate non-metric multidimensional scaling visualizations using Bray-Curtis dissimilarities of algal symbiont community per species, and for the effects of site and treatment. To investigate the effect of treatment in the symbiont diversity, chao1, representing species richness, was calculated in R using the package *iNEXT* (*98*) and the function *ggINEXT*. The venn diagram was created using the package g*gVennDiagram* (*99*).

To assess the association between the numbers of samples that survived each treatment and the strategy they follow (AKP and ABH) we performed a Fisher Exact test in R.

Color for the figures 1b and 3a, b were generated using the package *MetBrew*

## Acknowledgments

We thank the Coral Resilience Lab (the legacy of Ruth Gates) and ToBo Lab who assisted with collections and laboratory work (special thanks to Becca Kitchen, Emily Conklin and Evan Barba Freel). We thank also Joan Roughgarden for providing valuable feedback on the interpretation of her holobiont model.

This is SOEST contribution number xxx and HIMB contribution number xxx.

Our studies were conducted under permits #14-1702-1 to 14-1702-5 from the University of Hawai’i Institutional Animal Care and Use Committee (IACUC), and oversight from the Animal Welfare Regulatory Compliance Office. Coral collections were made under Hawai‘i Department of Land and Natural Resources permits SAP 2015-17 and SAP 2016-69 issued to the Hawai‘i Institute of Marine Biology.

## Author contributions

Conceptualization and design: MRS, RJT

Molecular work: MRS, CPJ

Analysis and interpretation: MRS, JB, CD, RJT

Funding: MRS, CD,CPJ, RJT

Writing—original draft: MRS, CD, RJT

Writing—review & editing: MRS, JB, CPJ, CD, RJT

## Data and materials availability

All data are available in the main text or the Supplementary Materials. Symportal data will be submitted to Zenodo and accession number will be made publicly available upon manuscript acceptance.

