## Supplemental material for "Stony Coral Symbioses Show Variable Responses to Future Ocean Conditions"

Mariana Rocha de Souza* *et al.*

### Supplementary Text

Genetic diversity among experimental corals

Based on previous work with these species, we expected clonal replication among the sampled colonies to be on the order of about 1% or less (*100-103*) and for clones to be spatially aggregated in the field. We intentionally spread collections at each site across a broad area (at least 5 m apart among conspecifics) to avoid sampling adjacent colonies to minimize the risk of sampling clone mates (*103-105*). Given our low exclusion power from a few markers and consideration of possible marking error, this is a highly conservative test. Consistent with expectations, there were only two individuals flagged as potential clones in *P. compressa*. This species showed a single potential clone pair that were collected from the same geographic location but, these two potential clonal colonies exhibited different coloration and morphology, making it exceedingly unlikely that any of the corals were clonally derived.

Although the initial design was balanced, the number of genets who survived in each treatment through the end of the experiment varied per species (**Supplementary Table 2**). The ratio of the number of fragments per genotype in each tank treatment is close to 1, showing that there is high genetic diversity in each treatment, and confirming that results are not inflated by either the inclusion of clones or the survival of fragments from a few resilient genotypes.

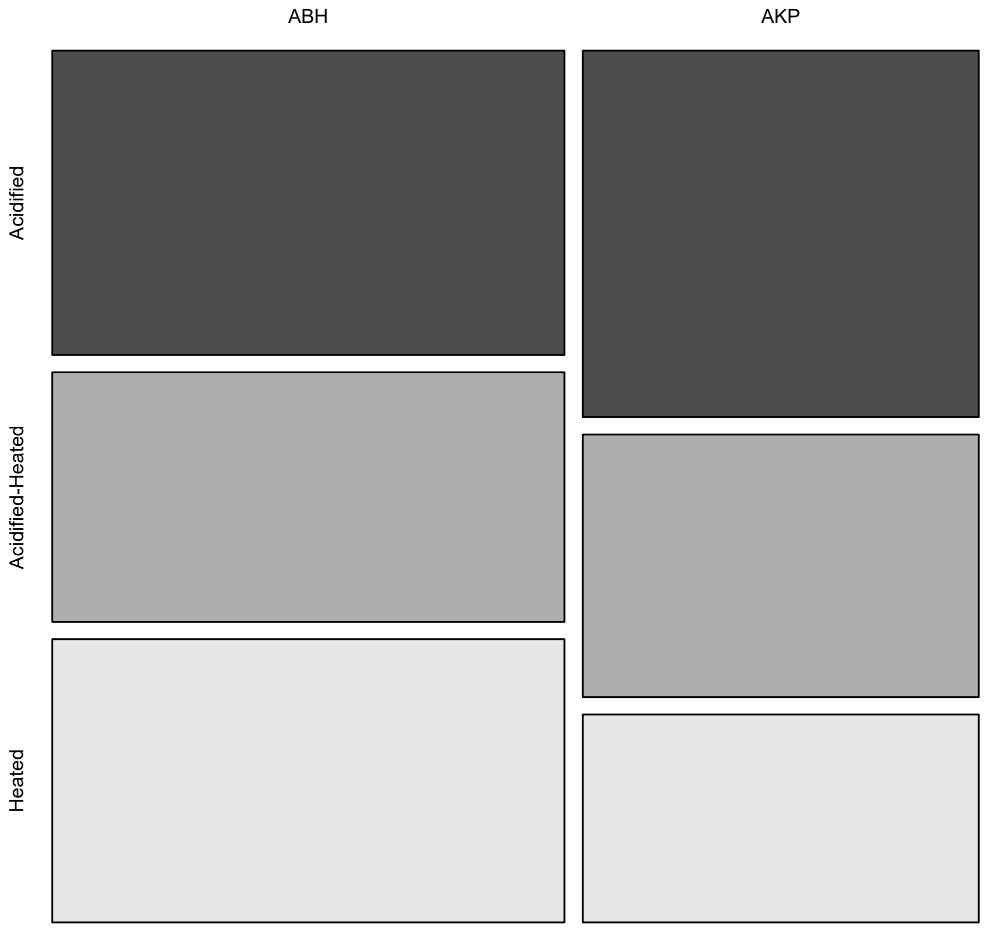

**Fig. S1.**
Mosaic plot showing the number of corals that survived across treatments, categorized by their symbiont strategy. Fisher test p= 0.16. The Adaptive Bleaching Hypothesis (ABH) is represented by a negative Δq, indicating symbiont selection following a stress event. In contrast, the Anna Karenina Principle (AKP) is represented by a positive Δq, reflecting an increase in symbiont diversity after a stress treatment.

| Collection site | Coordinates | Mean annual SST[a] (°C) | Mean summertime SST (°C) |
|---|---|---|---|
| HIMB | 21.43417 N, 157.78634 W | 26.14 ± 2.00 | 28.44 ± 1.04 |
| Sampan | 21.45239 N, 157.79487 W | 26.14 ± 2.00 | 28.44 ± 1.04 |
| Magic Island | 21.28754 N, 157.85037 W | 26.19 ± 1.35 | 27.68 ± 0.75 |
| Electric Beach | 21.35321 N, 158.13129 W | 26.34 ± 1.40 | 27.85 ± 0.75 |
| Hale‘iwa | 21.59252 N, 158.11034 W | 25.80 ± 1.19 | 27.06 ± 0.79 |
| Waimānalo | 21.32629 N, 157.67460 W | 26.00 ± 1.41 | 27.55 ± 0.89 |

**Table S1.**

Summary of environmental conditions at the six coral collection sites surrounding O‘ahu, HI. Table adapted from Price et al (2021).

| **Species** | **Initial collection** | **Acidified-Heated** | **Heated** | **Acidified** | **Control** |
|---|---|---|---|---|---|
| *Montipora capitata* | 30 | 9 (5) | 10 (6) | 29 (24) | 21 (19) |
| *Montipora flabellata* | 22 | 23 (15) | 22 (17) | 24 (16) | 18 (15) |
| *Montipora patula* | 30 | 21 (19) | 23 (17) | 27 (22) | 20 (17) |
| *Porites compressa* | 30 | 31 (25) | 27 (22) | 30 (25) | 28 (22) |
| *Porites evermanni* | 30 | 29 (21) | 28 (23) | 29 (23) | 28 (22) |
| *Porites lobata* | 30 | 25 (20) | 28 (24) | 28 (24) | 27 (23) |
| *Pocillopora acuta* | 30 | 9 (8) | 8 (8) | 18 (16) | 17 (15) |
| *Pocillopora meandrina* | 30 | 1 (1) | 1 (1) | 21 (18) | 12 (10) |
| **Total** | 232 | 148 | 147 | 206 | 171 |

**Table S2.**

Number of samples included in the study. Number of colonies (genets) at the initial collection which were then fragmented into treatments, and number of replicate clonal fragments (ramets) of each genetically unique coral colony (genet) from the initial collection that survived the four treatments. Numbers in parenthesis represent the number of genotypes found in each treatment.

| Treatments | Df | Sum Sq | F | $R^2$ | P |
|---|---|---|---|---|---|
| Treatments | 4 | 2.082 | 5.816 | 0.026 | 0.001* |
| Control vs Acidified treatment | 1 | 0.220 | 2.394 | 0.006 | 0.43 |
| Control vs Heated treatment | 1 | 0.6773 | 8.056 | 0.025 | 0.01* |
| Control vs Acidified-Heated treatment | 1 | 0.577 | 6.742 | 0.021 | 0.01* |
| Control vs Initial collection | 1 | 0.415 | 4.382 | 0.011 | 0.05* |
| Acidified vs Heated treatment | 1 | 0.767 | 8.808 | 0.024 | 0.01* |
| Acidified vs Acidified-Heated treatment | 1 | 0.653 | 7.371 | 0.021 | 0.01* |
| Acidified vs Initial collection | 1 | 0.116 | 1.202 | 0.003 | 1 |
| Heated treatment vs Acidified-Heated treatment | 1 | 0.034 | 0.439 | 0.001 | 1 |
| Heated treatment vs Initial collection | 1 | 0.967 | 10.730 | 0.029 | 0.01* |
| Acidified-Heated treatment vs Initial collection | 1 | 0.805 | 8.79 | 0.024 | 0.01* |

**Table S3.**

Pairwise PERMANOVA of the Symbiodiniaceae community in each fragment exposed to the different treatments. Significance is indicated with *

| Acidified heated | | Heated | | Acidified | |
|---|---|---|---|---|---|
| Symbiont | R2 | Symbiont | R2 | Symbiont | R2 |
| D4 | 0.697 | D4 | 0.739 | D4 | 0.538 |
| D1 | 0.771 | D1 | 0.833 | D1 | 0.598 |
| D6 | 0.695 | D6 | 0.822 | D6 | 0.566 |
| D1dg | 0.477 | D1dg | 0.524 | D1dg | 0.401 |
| C15 | 0.734 | C15 | 0.756 | C15 | 0.736 |
| C1b | 0.425 | C1b | 0.429 | C1b | 0.422 |
| C42.2 | 0.453 | C42.2 | 0.450 | C42.2 | 0.461 |
| C1 | 0.448 | C1 | 0.443 | C1 | 0.440 |
| C3 | 0.435 | C3 | 0.502 | C3 | 0.479 |

**Table S4.**

Highest biplot Symbiodiniaceae loadings in each stress treatment.

| | Chi square | Degree of freedom | P |
|---|---|---|---|
| Species | 3336.06 | 7 | $<2.2e^{-16}$ * |
| Treatment | 27.48 | 2 | $1.080e^{-0.6}$ * |
| Site | 55.90 | 5 | $8.512e^{-11}$* |

**Table S5.**

Analysis of deviance of the variables used in the experiment. Significance is indicated with *

| | LR Chisq | Df | q |
|---|---|---|---|
| Species | 62.270 | 7 | <0.001 * |
| Treatment | 2.595 | 2 | 0.273 |
| Site | 2.796 | 5 | 0.731 |
| Species: Treatment | 19.813 | 12 | 0.071 |
| Species: Site | 50.343 | 23 | 0.001 * |
| Treatment: Site | 9.544 | 10 | 0.481 |
| Species:Treatment: Site | 18.096 | 30 | 0.957 |

**Table S6.**

Analysis of Type II Deviance Table with the variables included in the study. Significance is indicated with *

### Supplementary Materials References
